# Micromolar fluoride contamination arising from glass NMR tubes and a simple solution for biomolecular applications

**DOI:** 10.1101/2024.02.12.579991

**Authors:** Khushboo Matwani, Jasmine Cornish, Erika Alden DeBenedictis, Gabriella T. Heller

## Abstract

Fluorine (^19^F) NMR is emerging as an invaluable analytical technique in chemistry, biochemistry, material science, and medicine, especially due to the inherent rarity of naturally occurring fluorine in biological, organic, and inorganic compounds. Thus, we were surprised to identify an unexpected peak in our ^19^F NMR spectra, corresponding to free fluoride, which appears to leach out from various types of new and unused glass NMR tubes over the course of several hours. We quantified this contaminant to be at micromolar concentrations for typical NMR sample volumes across multiple glass types and brands. We find that this artefact is undetectable for samples prepared in quartz NMR tubes within the timeframes of our experiments. We also observed that pre-soaking new glass NMR tubes combined with rinsing removes this contamination below micromolar levels. Given the increasing popularity of ^19^F NMR across a wide range of fields, the long collection times required for relaxation studies and samples of low concentrations, and the importance of avoiding contamination in all NMR experiments, we anticipate that our simple solution will be useful to biomolecular NMR spectroscopists.

## Main text

Over the past seven decades, fluorine (^19^F) NMR has emerged as an invaluable analytical technique for a wide range of scientific applications, including chemistry, biochemistry,^(1-6)^ material science,^(7,8)^ and medicine.^(9,10)^ A distinguishing feature of ^19^F NMR, in addition to the remarkable sensitivity of the ^19^F nucleus, is the inherent rarity of naturally occurring fluorine in biological, organic, and inorganic compounds. Biochemically, ^19^F probes can be artificially introduced into proteins via post-translational modifications,^(11)^ or via incorporation of fluorinated amino acids into growth media.^(12) 19^F atoms can also be included within small molecules: to date, approximately 50% of all marketed agrochemicals,^(13)^ and more than 30% of recent FDA-approved pharmaceuticals contain ^19^F.^(14-16)^ Consequently, background interference, which is a common concern in other NMR nuclei, should, in principle, be virtually non-existent in the context of ^19^F NMR. Thus, when performing some routine ^19^F NMR experiments of 5-fluoroindole (with an expected ^19^F chemical shift near -126 ppm) in phosphate buffered saline (PBS), we were surprised to detect a second sharp peak near -120 ppm (**Fig 1**). The chemical shift of this peak is in close proximity to and could easily be mistaken for peaks of biochemical and/or medicinal chemistry interest, for example ^19^F-labelled tryptophan residues (labelled with 5-fluoroindole) which often occur in range -123 to -126 ppm, ^(17)^ and several fluoro-aromatic compounds which often fall in the range between -60 and -172 ppm.^(18-20)^ To our surprise, the unexpected peak appeared in spectra measured of solutions containing buffer alone across several brands, types, and classes of new NMR 5 mm tubes (**Fig 2**), including Wilmad Precision (type I, class A), Wilmad High Throughput (type I, class B), and Norell Secure (type I, class B). The contamination was also present to varying degrees across different buffers (**Fig 3**), but undetectable within the measurement time for samples prepared in new quartz tubes (Wilmad), suggesting the source of the contamination is the glass tube itself (**Fig 2a**).

**Fig 1:**
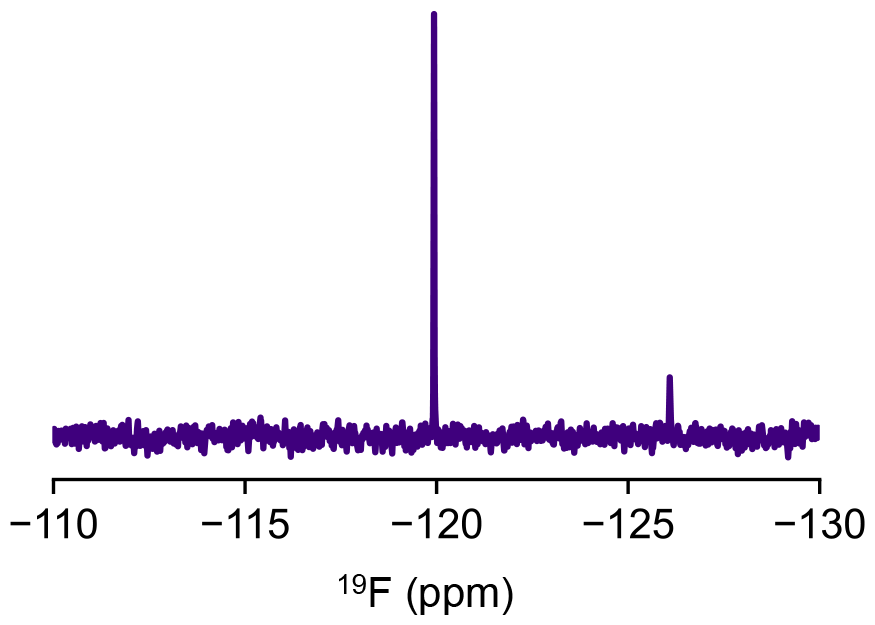
^19^F NMR spectrum of 20 μM 5-fluoroindole in PBS buffer, showing the expected peak for 5-fluoroindole near -126 ppm and an unexpected peak near -120 ppm.

**Fig 2:**
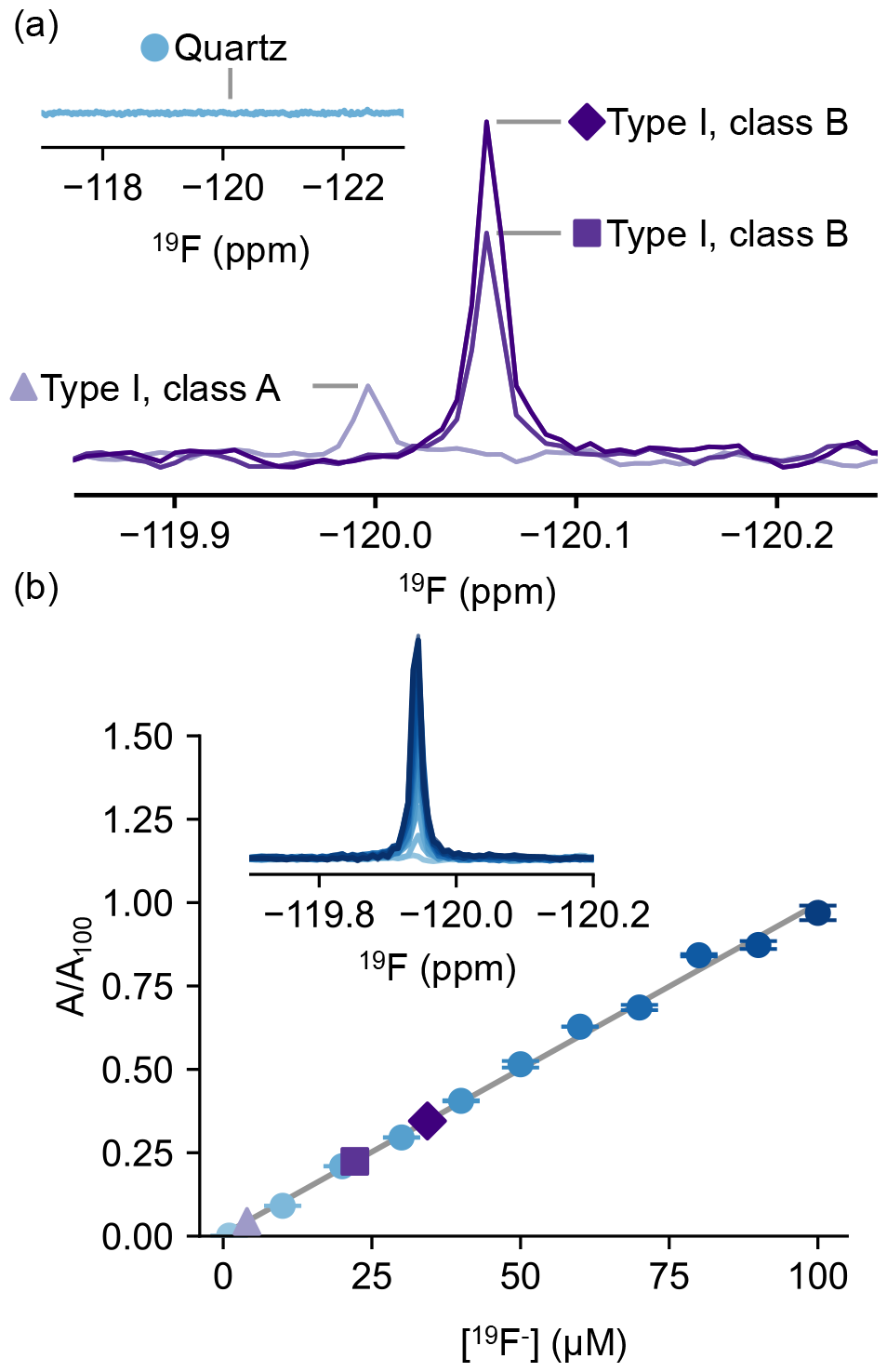
The unexpected peak near -120 ppm is observed in samples containing buffer alone in three types of glass NMR tubes, but not in quartz tubes. **(a)** ^19^F spectra of PBS incubated in various types of new, unused glass NMR tubes for 48 hours. Triangle: Wilmad Precision, type I, class A; square: Wilmad High Throughput, type I, class B; diamond: Norell Secure, type I, class B. Insert shows ^19^F spectra of PBS incubated in a new quartz tube for 48 hours. **(b)** Quantification of unknown fluoride concentrations in various glass tubes. Insert shows ^19^F spectra of known fluoride standards measured in quartz tubes. Normalised peak area is plotted as a function of fluoride concentration to determine fluoride quantities in various tube types. Triangle: Wilmad Precision, type I, class A (4 μM); square: Wilmad High Throughput, type I, class B (22 μM); diamond: Norell Secure, type I, class B (34 μM). Error bars represent one SD.

**Fig 3:**
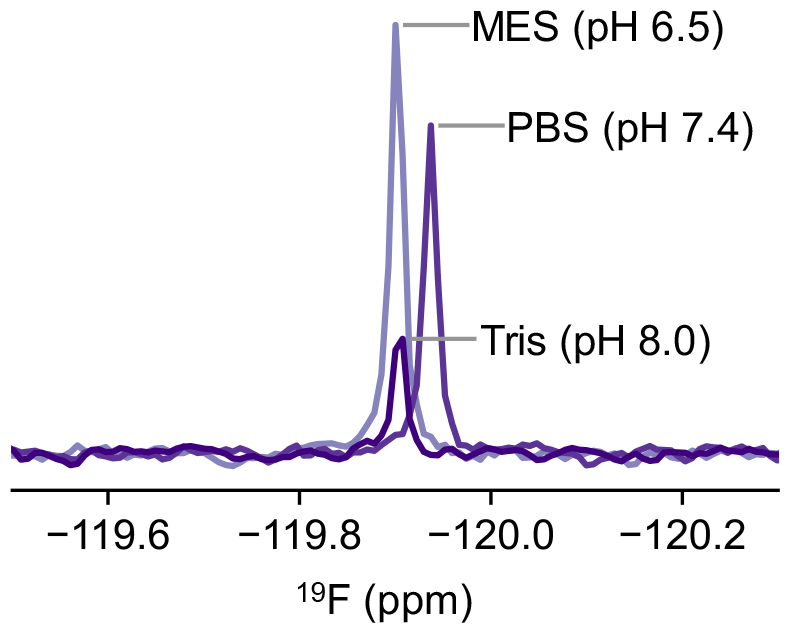
The unexpected peak from fluoride is observed in various buffers of diverse pH values after incubation in glass tubes (type I, class B) for 48 hours.

As the chemical shift of the contaminant peak is close to the reported value for fluoride, we prepared control samples of sodium fluoride in the same buffer in pre-rinsed quartz tubes and observed peaks of a similar nature to the artefact (e.g. chemical shift, peak widths). Using sodium fluoride standards ranging from 1 to 100 μM in pre-rinsed quartz tubes, we quantified the concentrations of free fluoride leaching from the various glass tubes (**Fig 2b**). We quantified free fluoride to be at micromolar concentrations after 48 hours of incubation for all glass tubes; we approximate the concentrations of free fluoride to be 34 μM for Norell Secure (type I, class B) tubes, 22 μM for Wilmad High Throughput (type I, class B), and 4 μM for Wilmad Precision (type I, class A) tubes.

Quartz has a modular molecular structure formed of highly regular tetrahedral SiO_2_ units, with minimal impurities,^(21,22)^ potentially explaining why we were unable to detect fluoride in quartz tubes. Borosilicate glass on the other hand, from which typical glass NMR tubes are made, is comprised of an amorphous structure consisting of boron atoms in a tetrahedral or trigonal conformation with silicon and bridging oxygen atoms.^(23)^ Fluorides are introduced during the refinement process, acting as a nucleating agent which increases the durability of glass.^(24,25)^ Additives, such as fluorides, further modify the glass network, increasing the pore size of the network formation.^(26)^ Fluoride,^(27-29)^ and other ions,^(30-32)^ have been shown to leach from different glass types over time. The dissolution process can take place over the span of hours and days as demonstrated in other studies.^(31)^ To probe the time dependence of fluoride leaching, we monitored fluoride signal intensities in ^19^F NMR spectra. Within an hour of placing buffer in a new Norell Secure (type I, class B) NMR tube, we collected spectra approximately every hour for 92 hours (**Fig 4**). We observed a steep increase within the first 10 hours, and with the fluoride content eventually tapering off to a plateau value of approximately 31 μM (**Fig 4**).

**Fig 4:**
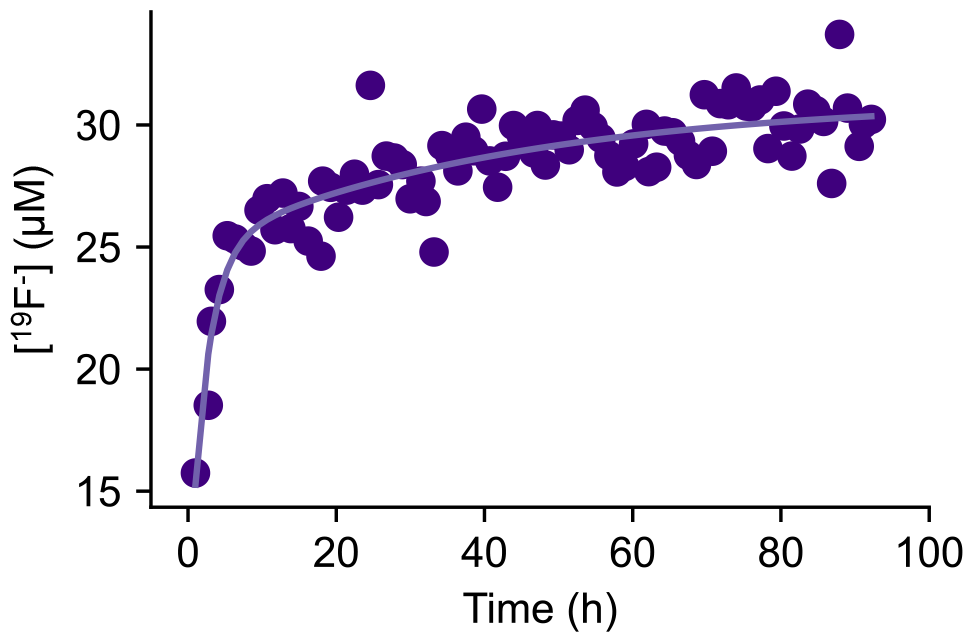
The release of free fluoride over time. A freshly made sample of PBS was incubated in a new, unused glass NMR tube (type I, class B) and data was collected approximately every hour. The integral of the growing peak over time was converted to a concentration based on the linear fit in **Fig 2**. The plateau value was determined to be 31 μM (see Methods).

Biomolecular NMR experiments, particularly relaxation studies or measurements of samples containing low concentrations, can often take several hours, and commonly samples are stored for several days or longer between measurements. While using quartz tubes is one simple way to minimise fluoride contamination, quartz tubes can be costly (currently ranging from $48 USD to $68 USD per tube) as compared to high-field glass tubes (currently ranging from $9 USD to $54 USD per tube). We therefore wondered whether extended incubation and subsequent rinsing of the tube could remove the fluoride contamination. To test this, we incubated a Norell Standard (type I, class B) tube in PBS for 48 hours, and as we predicted, significant leaching of fluoride ions was observed. We then rinsed the tube twice (once with water and once with buffer) and recollected the ^19^F NMR spectrum. The absence of any peak near -120 ppm suggests that the fluoride level was below 1 μM (**Fig 5**).

**Fig 5:**
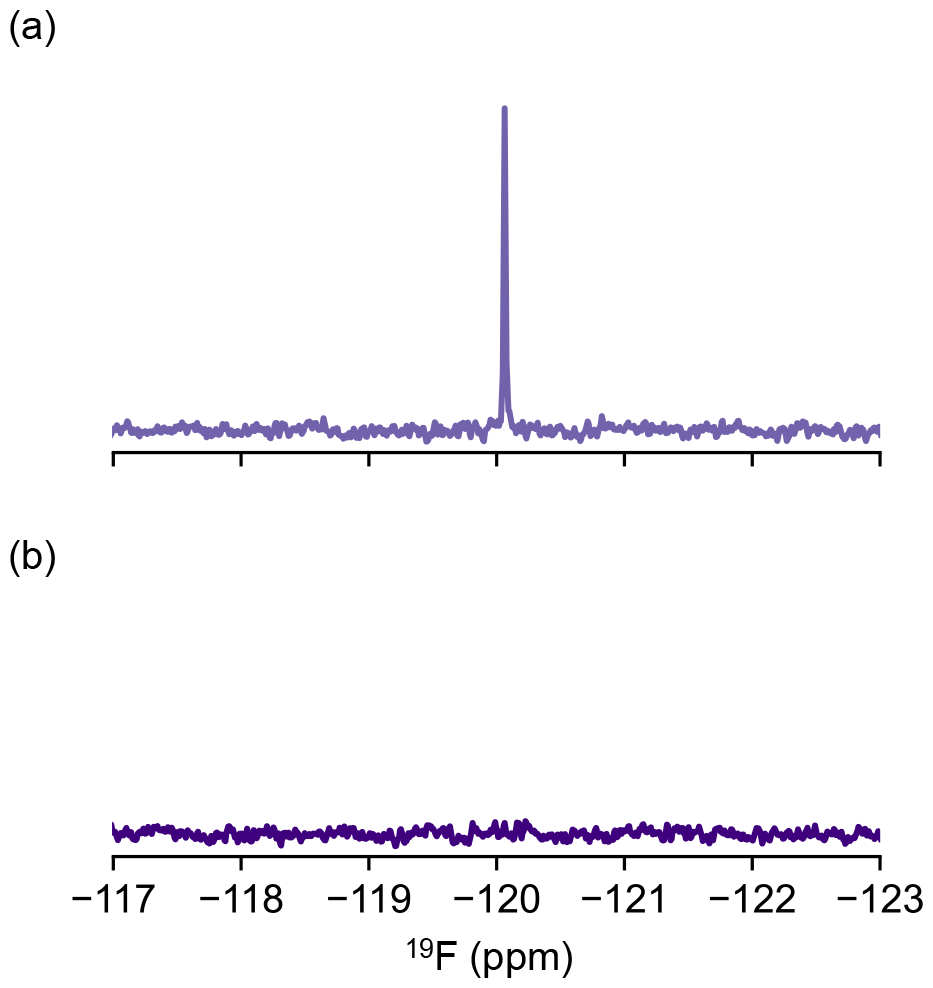
Pre-soaking NMR tubes with buffer is an easy strategy to eliminate the fluoride contaminant from glass. **(a)** ^19^F NMR spectrum collected of PBS buffer in a new glass NMR tube (type I, class B) which had been incubated with buffer for 48 hours. **(b)** The sample from (a) was removed from the NMR tube, rinsed once with water and once with buffer before being incubated again with PBS. The ^19^F NMR spectrum collected after the rinse suggests the fluoride concentration is below 1 μM.

Fluoride leaching from glass has been previously reported for other glass surfaces,^(27-29)^ and this is particularly relevant in the context of typical NMR tubes, as they have a relatively high surface area/volume ratio. Here, we report that various types and brands of new, unused glass NMR tubes can lead to micromolar contaminations of fluoride, comparable to the concentrations used in many biochemical and drug-discovery experiments, which can significantly complicate NMR analyses. In the context of biochemistry, fluoride has been shown to inhibit enzymes such as enolases^(33)^ and hydrolases,^(34)^ bind proteins, and is essential for the function of nucleic acid structures like riboswitches.^(35)^ For example, we observed that free fluoride, at the contamination concentrations reported herein, binds to various proteins including DNase (**Fig 6a**), bovine serum albumin^(36)^ (BSA, **Fig 6b**), and the intrinsically disordered domains 2 and 3 of the non-structural protein 5A (NS5A-D2D3, **Fig 6c**) from hepatitis C virus, indicated by a ^19^F chemical shift perturbation of the fluoride ion.

**Fig 6:**
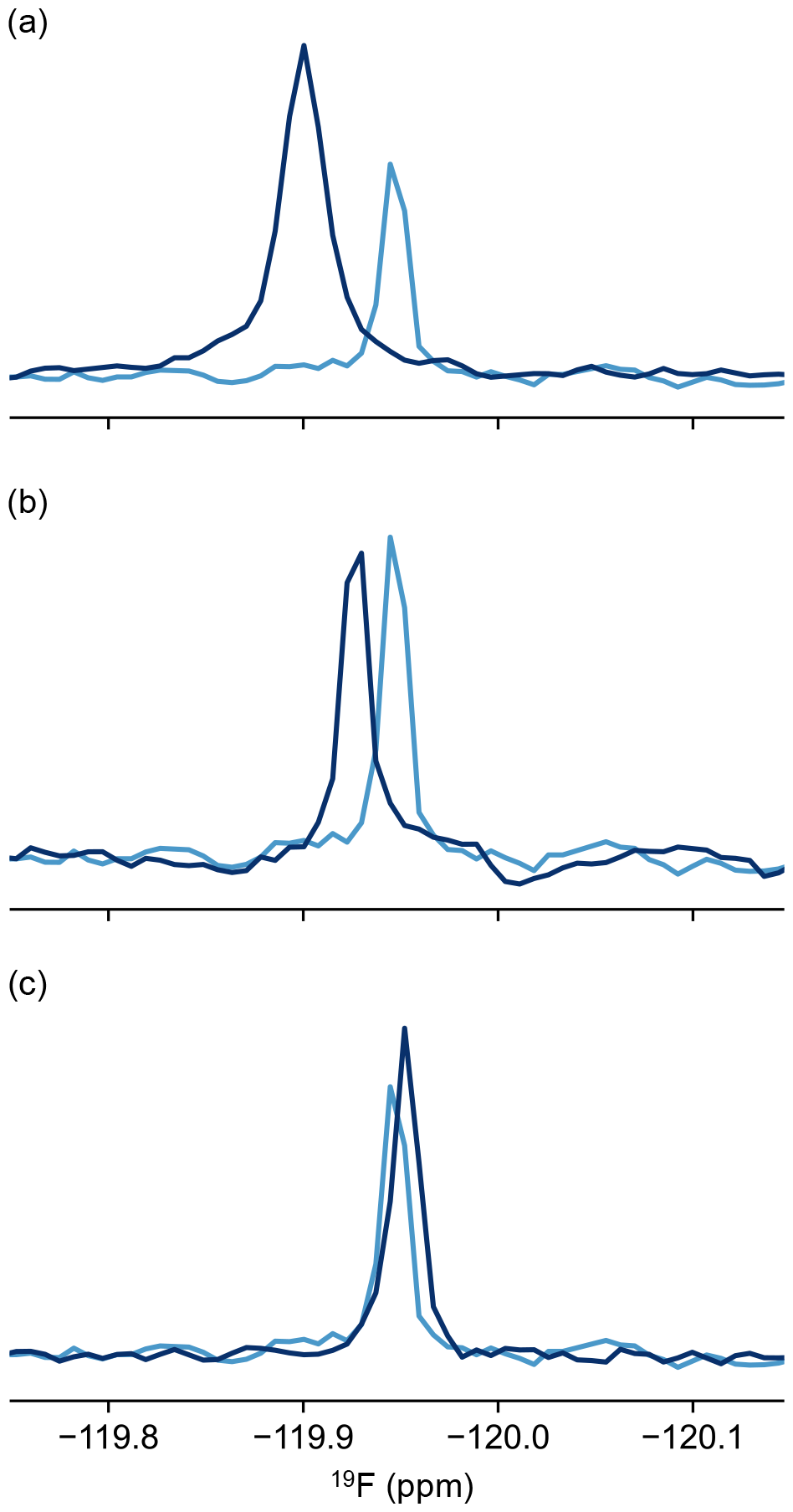
Chemical shift perturbations of fluoride suggest that the ion binds various proteins. ^19^F NMR spectra of a 20 μM sodium fluoride sample in the absence (light blue) and presence of various proteins (dark blue) including 100 μM DNase **(a)**, 200 μM BSA **(b)**, and 107 μM NS5A D2-D3 **(c)**.

Artefacts arising from long incubation times are particularly relevant for dynamic studies (e.g. Carr-Purcell-Meiboom-Gill and Chemical Exchange Saturation Transfer experiments)^(37)^ which can last from hours to days. In the context of ^19^F drug screening, the fluoride signal may be confused for signals arising from small molecules (e.g. fluoroaromatics), contaminants, impurities, or may further complicate interaction studies.^(38)^ While our study is limited to only a few sample conditions, we anticipate that this simple, inexpensive solution for removing fluoride contaminations by pre-soaking tubes will improve the quality of not only ^19^F spectra, but also of all biomolecular NMR data containing molecules which are sensitive to fluoride.

## Methods

### Sample preparation

Phosphate buffered saline (PBS) buffer (pH 7.4) was prepared by dissolving PBS tablets (Dulbecco A, Oxoid) in water according to the manufacturer’s instructions. 2-(N-morpholino)ethanesulfonic acid (MES) buffer was made at 25 mM with 140 mM NaCl and the pH was adjusted to 6.5. Tris(hydroxymethyl)aminomethane (tris) buffer was made at 25mM with 140 mM NaCl and the pH was adjusted to 8.0.

A recombinant construct of the D2 and D3 domains of HCV NS5A (NS5A-D2D3, residues 247-466) was expressed and purified as previously described.^(38)^ NS5A-D2D3 was prepared in PBS with 1 mM tris(2-carboxyethyl)phosphine (TCEP), pH 7.4. The protein was concentrated using an Amicon Ultra Centrifugal filter with a 3 kDa cutoff.

DNAse I (Roche) and BSA (Sigma-Aldrich) were made up to 1 mM in PBS with 1 mM TCEP, pH 7.4, and then diluted to 100 μM and 200 μM respectively.

5-fluoroindole (Sigma-Aldrich) was dissolved in DMSO-d_6_ at 1 M concentration and kept frozen at -20°C until use.

Samples of PBS buffer and 2% D_2_O were prepared to a final volume of 550 μL loaded into three different types of new 5mm NMR glass tubes: Norell Secure (type I, class B), Wilmad Thin Wall Precision (type I, class A), and Wilmad High Throughput (type I, class B) and incubated for over 48 hours. MES and tris buffers with 2% D_2_O were prepared to a final volume of 550 μL, loaded into new 5 mm Norell Standard NMR tubes (type I, class B) and incubated over 48 hours. As a control, PBS buffer was incubated for the same time period in a new quartz tube (Wilmad).

Sodium fluoride standards were prepared in PBS buffer (pH 7.4) at concentrations ranging from 1 μM to 100 μM and measured in pre-rinsed quartz tubes.

### NMR experiments

^19^F spectra were collected on a 18.8 T Bruker Avance III 800 MHz spectrometer equipped with a TCI cryoprobe at 298K. ^19^F spectra were acquired using the standard aring Bruker pulse sequence to minimise acoustic ringing. Spectra were obtained with 8,192 complex points, a spectral width of 45,455 Hz, a relaxation delay of 0.5 s, and an acquisition time of 0.09 s. 6,400 scans were collected per experiment, and the offset was set to -120.00 ppm. ^19^F chemical shifts were referenced with respect to trichlorofluoromethane (CFCl_3_). Data was processed and analysed using nmrPipe,^(39)^ and nmrGlue.^(40) 19^F peaks were fit to Lorentzian curves.

Peak integrals of known fluoride standards were calculated and plotted as a function of fluoride concentration. A linear fit of the data was used to calculate unknown fluoride concentrations in various glass tubes.

To investigate the kinetics of the leaching process, a sample was prepared as described above and added to a Norell Secure tube (type I, class B). Data collection began within one hour of placing the sample in the NMR tube. Peak integrals were fit to the following bi-exponential curve:

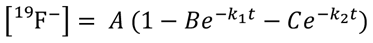

where *A* is the plateau of the curve, *t* represents time, and *B, C, k*_1_ and *k*_2_ are constants.

Free fluoride was removed from a Norell Standard NMR tube (type I, class B) by pre-soaking for 48 hours with PBS buffer. The tube was then rinsed twice, once with water and once with PBS before a fresh sample was added, and a spectrum was collected.

## Acknowledgements

The authors acknowledge Prof. D. Flemming Hansen, Dr Geoff Kelly, and Prof. Snezana Djordjevic for helpful discussions. GTH was supported by a BBSRC Discovery Fellowship (BB/X009955/1). JC was supported by a Steel Perlot Early Investigator Grant. The BBSRC (BB/R000255/1), Wellcome Trust (ref101569/z/13/z), and the EPSRC are acknowledged for supporting the NMR facility at University College London. Access to ultra-high field NMR spectrometers was supported by the Francis Crick Institute through provision of access to the MRC Biomedical NMR Centre. The Francis Crick Institute receives its core funding from Cancer Research UK (FC001029), the UK Mesdical Research Council (FC001029), and the Wellcome Trust (FC001029). This paper was typeset with the bioRxiv word template by @Chrelli: www.github.com/chrelli/bioRxiv-word-template.

## Competing interest statement

The authors declare no competing interests.

## Notes

### Competing Interest Statement

The authors have declared no competing interest.

